# Incorporating structural knowledge into unsupervised deep learning for two-photon imaging data

**DOI:** 10.1101/2021.05.18.443587

**Authors:** Florian Eichin, Maren Hackenberg, Caroline Broichhagen, Antje Kilias, Jan Schmoranzer, Marlene Bartos, Harald Binder

## Abstract

Live imaging techniques, such as two-photon imaging, promise novel insights into cellular activity patterns at a high spatio-temporal resolution. While current deep learning approaches typically focus on specific supervised tasks in the analysis of such data, we investigate how structural knowledge can be incorporated into an unsupervised generative deep learning model directly at the level of the video frames. We exemplify the proposed approach with two-photon imaging data from hippocampal CA1 neurons in mice, where we account for spatial structure with convolutional neural network components, disentangle the neural activity of interest from the neuropil background signal with separate foreground and background encoders and model gradual temporal changes by imposing smoothness constraints. Taken together, our results illustrate how such architecture choices facilitate a modeling approach that combines the flexibility of deep learning with the benefits of domain knowledge, providing an interpretable, purely image-based model of activity signals from live imaging data.

**Teaser sentence:** Using a neural network architecture that reflects domain knowledge provides an interpretable model of live cell imaging data.

## 1 Introduction

In the last decade, deep learning-based approaches have been successfully adapted for the analysis of image data, rivaling and surpassing human performance in identifying structure from data. This encompasses both supervised tasks with ground truth information, such as image segmentation with the U-Net [1], and unsupervised tasks where structure can be uncovered without prior human labeling, e.g., by generative models such as variational autoencoders (VAEs) [2]. Recently, in particular such unsupervised approaches have been extended to incorporate structural knowledge on temporal patterns in the form of explicit models [3, 4].

In line with such ideas of a gradual transition from black-box deep learning towards more explicit models we want to illustrate approaches for incorporating structural knowledge into deep generative modeling of live cell imaging data. Besides convolutional neural networks, which take into account spatial structure, we propose an architecture for distinguishing between the image foreground, containing the biological signal of interest, and a background. In addition, knowledge about gradual temporal changes in the biological processes being observed is incorporated in the architecture.

Specifically, we explore how a generative deep learning model based on VAEs can be adapted to such live imaging data. Exemplarily, we consider data from an *in-vivo* two-photon imaging experiment and show how gradually incorporating constraints and structural assumptions that reflect the properties of the data can provide an interpretable general model of neural activity. Such a model is then not specifically adapted to a particular prediction task but can be flexibly used for various downstream analyses.

Two-photon calcium imaging, as an exemplary application to illustrate our approach, represents an invasive technique in the context of ‘neural decoding’ for recording activities of individual neurons over time. Here, two-photon microscopy is used to capture movies of a neuronal population that expresses a fluorescent calcium indicator and allows to visualize the increase in intracellular Ca2+-concentration that accompanies neuronal spiking activity [5]. More generally, ‘neural decoding’ refers to techniques that use brain signals to make predictions about behavior, perception, or cognitive state and are becoming increasingly important for neuroscientific research [6–9]. For example, calcium imaging techniques enable optical measurements of large neural populations at a high spatio-temporal resolution. The resulting calcium imaging movies thus facilitate insights into neural activity [5, 10, 11]. To realize their full potential, computational methods are needed to derive a model of neural activity from the indirect optical measurement of the Ca2+ indicator. Here, to analyze the resulting video data, the typical two-step analysis framework of cell population imaging is employed, where cells are identified through segmentation in a first step, and the signals of interest, e.g., temporal fluorescence traces or firing rates, are identified subsequently [9, 12–14]. Current approaches to derive an explicit generative model typically rely on simplifying assumptions of, e.g., a linear combination of additive signals, that allow to directly incorporate known structure, but limit the flexibility and versatility of the obtained data model [15, 16]. On the other hand, deep learning techniques provide a more flexible class of models and have recently been successfully applied to infer spike activity from calcium imaging data (e.g., [17–19]). Yet, as more opaque black-box approaches, they lack the interpretability of an explicit data model and have so far been mainly used on two-photon imaging data for supervised prediction tasks, requiring large amounts of labeled training data that is typically not available.

Additionally, these approaches often only provide one of several components in the entire data analysis workflow [14], where prior steps, such as motion correction, can considerably affect performance [20]. Yet, results showing that temporal information can improve segmentation [17] indicate that it might be beneficial to consider both steps simultaneously and perform modeling directly based on the temporal sequence of images without explicitly extracting traces [9]. Such an integrated modeling strategy can help to build a flexible and versatile model that is not limited to a specific task.

We hypothesize that unsupervised generative deep learning approaches, which provide a model of the underlying data generating distribution, can be useful to build such integrated models. Specifically, we consider VAEs for this task, as they can be trained in an unsupervised way and infer a lowdimensional latent representation of the central factors of variation underlying the data. This facilitates interpretability and has been shown to be useful for capturing and extracting patterns in the data in an explainable artificial intelligence (AI) approach [21]. While VAEs have been adapted to two-photon imaging data for the specific task of inferring neural spike rates from fluorescence traces [19], we aim to exemplify how a deep learning approach based on a VAE architecture can be adapted to provide a flexible and versatile model that is not restricted to a specific task, but tailored to the properties of two-photon imaging data.

This paper is structured as follows. We first give an overview of our proposed modeling approach of gradually incorporating increasing levels of structural information into a VAE architecture. Next, we present results of the models trained on a two-photon imaging dataset of hippocampal CA1 neurons and discuss our findings. We additionally provide an overview of the typical image processing workflow as well as an introduction to generative deep learning in general and VAEs in particular, and the convolutional architecture integrated in our models. Finally, we give details on the two-photon dataset and the proposed modeling approaches and their implementation.

The source code of the implementation and an accompanying Jupyter notebook illustrating an exemplary application of the approach is available at https://gitlab.imbi.uni-freiburg.de/maren/incorporating-structural-knowledge-into-unsupervised-deep-learning-for-two-photon-imaging-data.

## 2 Results

### 2.1 Adapting a VAE-based model to the properties of two-photon imaging data by incorporating different amounts of structural knowledge

We employ a VAE as an unsupervised generative deep learning approach to model neuronal activity based on two-photon imaging movies. Specifically, we consider a sequence of individual frames from such a video as input data and propose an approach to tackle the challenging modeling task of simultaneously capturing the foreground signal with the distinct cellular activity and the coarse-grained, blurry background signal, while accounting for the temporal correlation between neighboring frames. Starting with a simple model architecture, we gradually increase the level of additional structural information integrated into the model. Thus, we explore how different encoded structural properties affect the activity patterns and the identified latent structure.

Figure 1 provides an overview of this framework, representing steps from a black-box model towards a more explicit model tailored to the data: In a first step, we train a fully-connected vanilla VAE on the dataset, as described in Section 4.3, where each image frame is treated as a separate observational unit. In a second step, we employ a convolutional architecture for the VAE encoder and decoder to account for the characteristics of the image data (see Section 4.4). Next, we tackle the mixed-source activity by building separate encoders and decoders for the foreground and background, respectively, that share a joint latent space. In a final model extension, we consider latent kernel smoothing to account for the temporal correlation across frames, thus incorporating a temporal smoothness constraint in the latent space.

**Figure 1:**
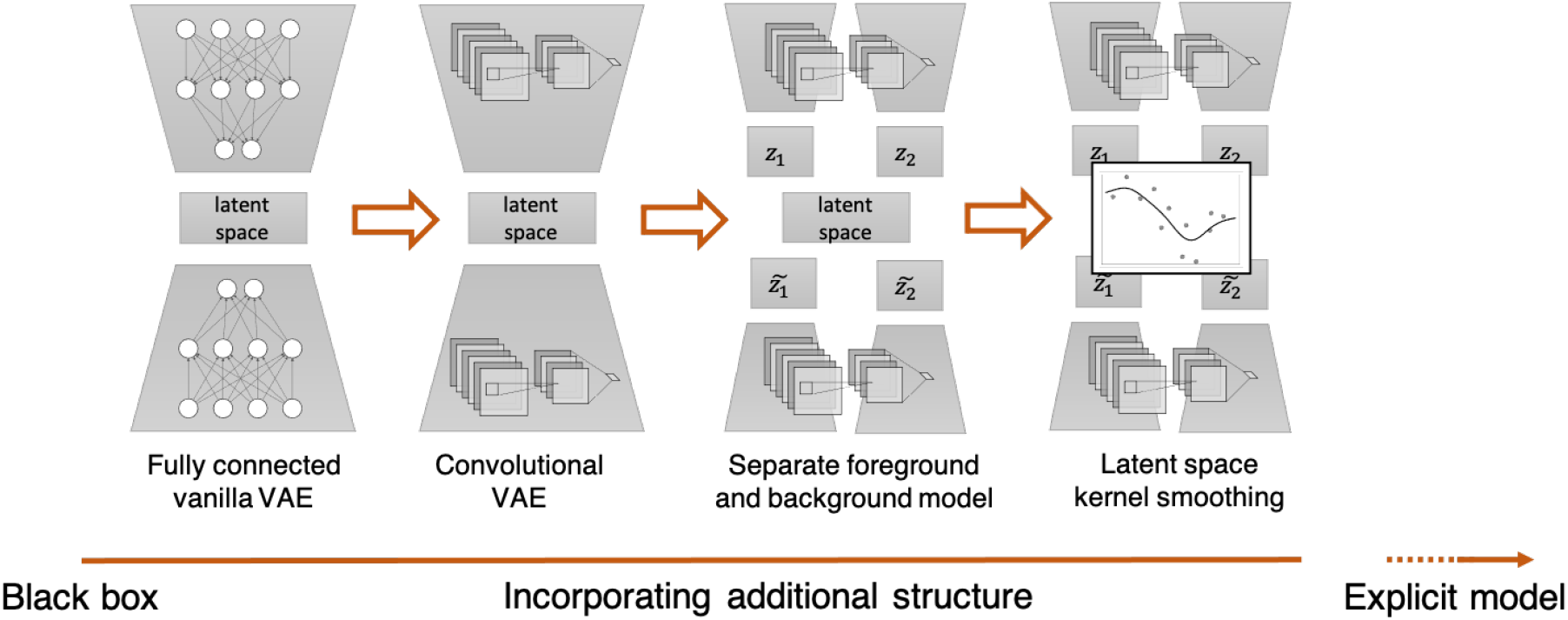
Overview of deep generative model architectures with increasing amounts of domain knowledge incorporated. Layers of structural information can be gradually combined with a fully connected vanilla VAE model (1) to enable a step-wise transition from a black-box deep learning model towards a more explicit modeling approach that takes into account problem structure by adding a convolutional architecture (2), separate foreground and background encodings (3), and latent kernel smoothing (4).

The recorded two-photon video frame images are characterized by a clear distinction (at least for the human observer) between foreground signal, showing the somatic calcium increase during firing of the neuron over time that we aim to capture with our modeling approach, and background, consisting primarily of more coarse-grained, noisy neuropil signal (for details on the data, see Section 4.7). Hence, we explicitly model the dataset of images as a combination of foreground and background signal that we seek to disentangle in the latent space of our VAE model.

To achieve this, we introduce separate VAE models for the foreground and background with a distinct latent representation of the foreground and background respectively. We realize these models as convolutional neural networks (CNNs) to account for the spatial structure of the image: In a CNN layer, a filter weight matrix is convolved (i.e., multiplied element-wise and summed up) with different patches of the image, like a window sliding over the image with its size given by the size of the filter matrix (for details, see Section 4.4). This accommodates central features of image data, such as distinctive local motifs, e.g., groups of active neurons, that can appear in any part of the image. We employ CNNS with different filter sizes for the foreground and background model to accommodate both sharp and local foreground signal and blurry, large-spanning background signal. To obtain a joint model of the entire image including both foreground and background, we integrate the latent representations of both VAE models into a joint latent space. The resulting *dual-target VAE* model is trained by optimizing an adapted version of the evidence lower bound, a lower bound on the data likelihood (for details, see Section 4.5 and the accompanying Jupyter notebook).

Since the images correspond to frames in a video, subsequent images exhibit similar cellular activity patterns and are thus strongly temporally correlated. We explicitly incorporate this spatio-temporal information as a structural property of the data into our model for constraining the latent representation. This will allow to model smoother developments and thus more accurately reflect underlying activity patterns in the data. Specifically, we propose kernel smoothing in the latent space and train the model on neighborhoods of adjacent images, i.e., subsequent frames in the video sequence. For each frame, we consider a neighborhood of a fixed size around the frame and obtain the latent representation of this frame as a weighted sum of the latent representations of all frames in the neighborhood, where the weights are given by a kernel that takes into account distance between frames (see Section 4.6). In this way, we impose a smoothness constraint on the latent space in that each frame’s representation is also influenced by the representations of neighboring frames. This facilitates to capture gradual temporal changes in activity patterns. We additionally compare this approach to a similar smoothing step performed at the level of the loss function, i.e., for each frame, we take the weighted average of all loss values of the frames in its neighborhood as the loss.

In the following, we compare (1) a VAE with a CNN architecture, (2) a dual-target VAE, and (3) a dual-target VAE with kernel smoothing.

### 2.2 Convolutional VAE

First, we evaluate the performance of the baseline CNN approach against a standard VAE with a fully-connected ANN encoder and decoder. With this fully-connected approach, we were not able to achieve dynamic reconstruction of the foreground and thus omit the results. In contrast, Figure 2 exemplarily shows that incorporating a CNN architecture in the VAE encoder and decoder allows for good reconstruction results of prominent foreground cells. This implies that the advantages of CNNs over a standard fully-connected neural network structure also apply to frames from two-photon imaging videos. Modeling of theses images hence benefits from taking into account local motives and patterns as well as their spatial invariance across the entire image, the main characteristics motivating the CNN architecture.

**Figure 2:**
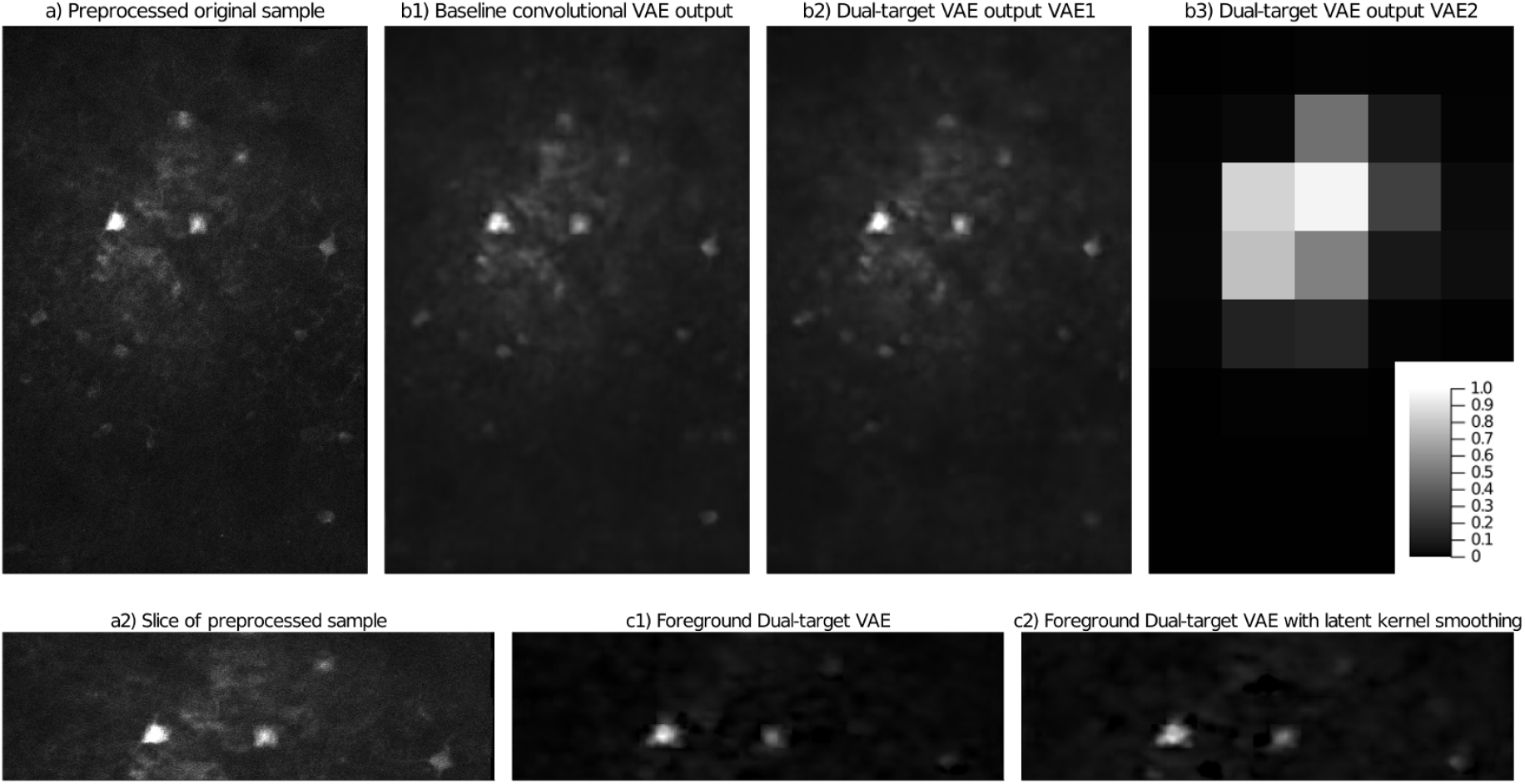
Comparison of reconstructions from a convolutional VAE, a dual-target VAE and a dual-target VAE with kernel smoothing. Top row: Preprocessed sample frame from the data (a) and its reconstructions of the convolutional baseline VAE approach (b1) and of dual-target VAE (b2) respectively. Bottom row: Slice of the same sample (a2) with corresponding reconstructions of only the foreground of dual-target VAE (c1) and dual-target VAE with latent kernel smoothing (c2).

Yet, the signal of smaller cells with Calcium-transients of shorter duration and smaller amplitudes is poorly reconstructed by the model in most of these smaller cells with lower firing intensity in Figure 2. This can be more generally observed over most other frames in the video, where the activity of the less prominent cells is reconstructed with low accuracy. Also, when observed over the course of contiguous frames, rapid alternations in cell activity, which are not present within the original data, are visible in the output underlining the problems of the model with this kind of signal. The noise in the original sample is not retained in the output, thus creating a slightly more blurred but denoised version of the input, which might be useful, e.g., for visual presentation.

### 2.3 Dual-target VAE

Next, we investigated the separation of foreground and background activity within the latent space of the convolutional baseline approach. To this end, we decoded one-hot configurations of all dimensions of the latent space (i.e., vectors with a value of 0 in all but one dimension) and observed the amount of foreground and background present in the output. In Figure 3, we display the outputs of the configurations with the most and least foreground activity. Even in dimensions with low foreground activity, at least some foreground cell activity is present and thus, this approach does not allow to fully disentangle fore- and background signal.

**Figure 3:**
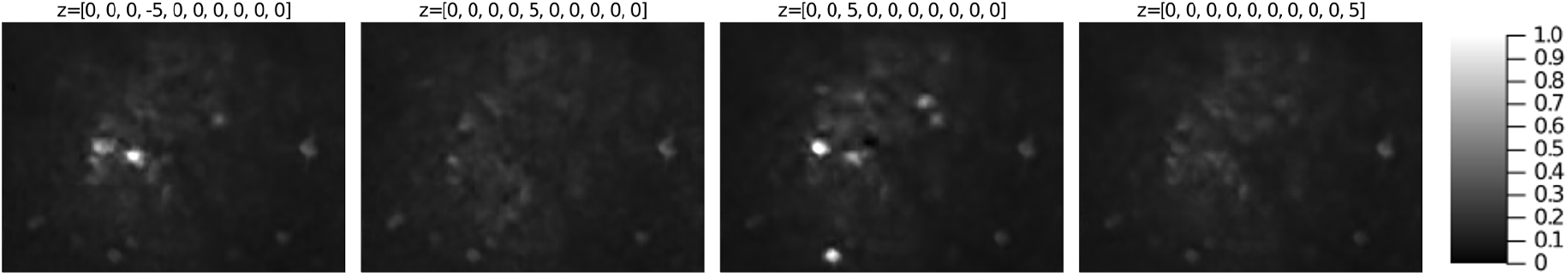
Exploration of the latent space dimensions with most and least foreground activity for a convolutional VAE and a dual-target VAE. Exemplary exploration plots (sliced) of foreground and background model of the baseline convolutional VAE model (left) and the dual-target VAE (right). The configuration of the latent space that was used to create the plot is annotated on top. Dimensions of the baseline model are chosen according to visual inspection for most and least foreground activity respectively.

On the other hand, the dual-target VAE showed no foreground activity within the two latent dimensions of the background VAE model and thus achieved better separation. For any given image, we can thus set **z**_1_ = 0 and decode the given configuration to an image of only the background activity. Subsequently, this background can be subtracted element-wise from the reconstructed image and we obtain an image of the foreground activity. Figure 2 exemplarily shows the reconstructed foreground of the given frame. The depicted frame is representative of the rest of the data, where static and dynamic parts of the background are eliminated with high accuracy and the dominant foreground cell activity is retained in the output. Yet there is still a tendency of the model to weaken or omit the activity of smaller, less active cells in the foreground.

### 2.4 Dual-target VAE with latent kernel smoothing

Finally, we evaluate the impact of kernel smoothing on the model performance by comparing the dual-target VAE with kernel smoothing on the latent representation and to the dual-target VAE without temporal smoothing (Figure 2): Comparing the foreground activity (bottom row) show that especially low-activity cells are more prominent in the reconstruction of the approach with latent kernel smoothing. Effectively, calculating the kernel weighted average of latent representations across a neighborhood of subsequent frames corresponds to imposing a smoothness constraint on the latent space. To further investigate whether this smoothness constraint can help to capture the temporal correlations between subsequent frames, we considered sequences of frames with a distance of two, i.e., used every second frame in the original video sequence for building the neighborhood. This means that the first, second and third frame in the neighborhood used for training the model correspond to the first, third and fifth frame in the respective part of the original video (see Figure 4). Specifically, we compared the differences between two adjacent frames in the sequence (i.e., between every second frame in the original video) when reconstructed with the dual-target VAE without kernel smoothing (Figure 4, b)) versus the dual-target VAE with latent kernel smoothing (Figure 4, c)). Here, we can observe that the differences between frames reconstructed based on a kernel smoothed latent representation are less pronounced than without the smoothing step.

**Figure 4:**
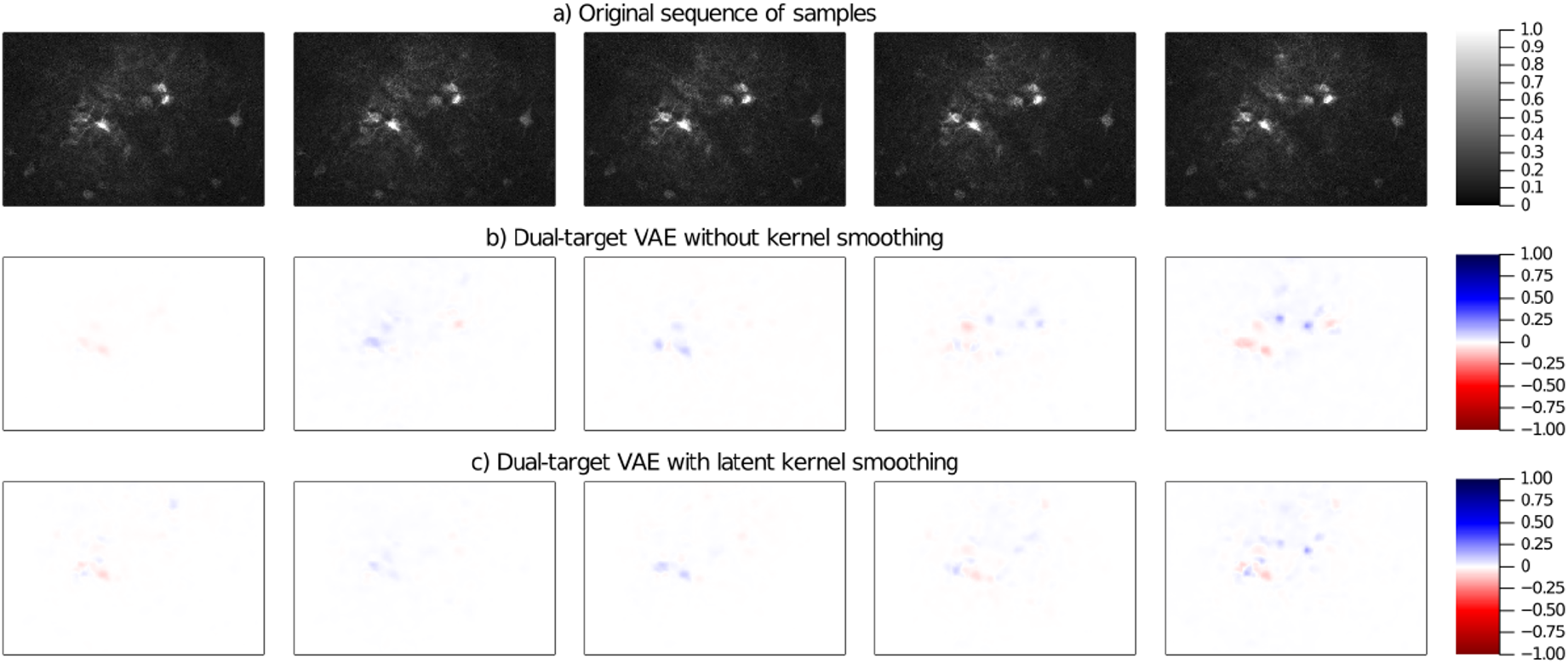
Comparison of differences between model reconstructions of adjacent frames with and without latent kernel smoothing. Exemplary sequence of frames and the differences between the model reconstructions of adjacent frames with and without latent kernel smoothing. a) Original exemplary sequence of samples with a distance of two, i.e., adjacent frames in the plot correspond to every second frame in the original video. b) Differences between the reconstructions of the frames shown in a) obtained from training a dual-target VAE without kernel smoothing. c) Differences between the reconstructions of the frames shown in a) obtained from training a dual-target VAE with kernel smoothing. The color coding describes the sign of the difference (blue: positive, red: negative), while the color intensity encodes the absolute magnitude.

We more generally observed a tendency towards higher intensity and precision of the reconstructed foreground activity with fewer flickering artifacts, indicating that our approach can indeed provide a smoother reconstruction of frames across time that reflects the underlying time-dependent biological process.

Additionally, we compare the kernel smoothing in the latent space with a dual-target VAE trained on neighborhoods with a smoothing step performed at the level of the loss values, as in 4, Section 4.6, i.e., using for each frame its individual latent representation, but employing a kernel weighted average across the loss values of all frames in a neighborhood as training objective. Here, the improvements described above are observed to a lesser extent (results not shown), implying that constrains should be imposed directly on the latent representation, which encodes the central structure underlying the data and the reconstruction.

## 3 Discussion and conclusions

Deep learning has been shown to be a useful tool for analyzing biomedical imaging data across a wide range of tasks. Specifically, such approaches have been developed for neural decoding [9] and are increasingly becoming a part of analysis workflows for live cell imaging data, such as for two-photon imaging [14, 17, 18]. Here, automatic extraction and modeling of neural activity from individual cells over time and in large cell populations is a crucial step towards a better understanding of cellular activity. This can ultimately be linked to phenotype data to facilitate biological insight. While current approaches are dominated by supervised models that require large amounts of labeled training data to learn a specific classification of prediction task, generative models infer a model of the underlying data distribution. Explicit data models that describe cellular activity as, e.g., linear combination of additive signals, are less opaque than deep learning approaches, but limited in their flexibility due to simplifying modeling assumptions. Ideally, a modeling approach should provide a combination of both, i.e., an interpretable model that allows to encode known structural properties of the data while being as flexible and versatile as possible. We have thus investigated how structural knowledge can be incorporated into an unsupervised generative deep learning approach. Specifically, we have employed VAEs, directly applied to the video frames from two-photon imaging data, for modeling neural activity in a latent representation. Here, one focus was on exploring how gradually incorporating structural components, which reflect key properties of two-photon imaging data, affect the latent representation, and thus facilitate insight into what the model has learned. Further, we exemplified to what extent a generative deep learning approach can be tailored to model signals activity in live cell imaging data.

Specifically, we have compared VAEs with different amounts of explicit structure incorporated into the model. While a standard fully-connected VAE did not permit to accurately reconstruct the cell images, employing convolutional layers, the *de facto* standard architecture for deep learning on image data, allowed to infer a latent representation based on which the input images could be reconstructed. Next, we have taken into account the typical mixed-source activity of two-photon imaging videos, where the fluorescent traces of active neurons in the background should ideally be separated and deconvolved from the noisy neuropil background signal. To model this structure, we have proposed separate VAE encoders and decoders for the foreground and background of each image, while still learning a joint latent representation of the image. With this, we have been able to obtain a latent representation that can disentangle foreground and background signals, which was only possible by explicitly encouraging it with the distinct encodings. Finally, we have illustrated a straightforward approach for imposing a smoothness constraint on the latent space, to take into account the temporal correlation and smooth structure of activity traces in subsequent frames. Specifically, we have formed neighborhoods of adjacent images around each frame and obtained the latent representation of each frame as a smoothed average of the activity signal over the entire neighborhood. This approach allowed to retain weaker signals from cells in the foreground more frequently and more clearly, thus capturing the overall activity more accurately by exploiting the similarity of subsequent frames for modeling.

Still, it is difficult to assess the model performance objectively beyond visual inspection of reconstructed videos and images, and more rigorous quantitative criteria are needed to evaluate further model refinement. Another drawback of the model in its current form is the complexity, due to many parameters in the convolutional layers, resulting in a computationally rather extensive training procedure. Additionally, more sophisticated approaches for smoothness constraints could be considered to facilitate dynamic modeling of the latent representation over time. For example, differential equations could be incorporated in the latent space. A further extension would be to consider stochasticity, and to more explicitly model the spatial structure. Here, recent works suggest underlying low-dimensional spatio-temporal dynamics in two-photon-imaging data [32], which could potentially be captured in the latent space of a VAE model, while the spatio-temporal patterns could be modeled with an approach as in [33]. Another interesting direction for future research are methods for linking the inferred low-dimensional activity patterns, e.g., to phenotype data regarding the behavior of the animal. This could also benefit from incorporating structural assumptions on the spatio-temporal dynamics, as in [34].

In the present work, our focus was on illustrating how such a deep learning-based approach can be adapted to the specific properties of the data using two-photon imaging as an exemplary application, rather than a detailed comparative study of existing deep learning approaches for cell imaging data. We specifically highlighted how modeling can benefit from incorporating elements of more explicit data models, and investigated how such structural assumptions affect the patterns learned by the model.

More generally, our approach illustrates how generative deep learning approaches can be combined with various amounts of structural assumptions for greater interpretability, and how the learned representations can be inspected to assess the influence of these assumptions. It can be seen as an example of the currently ongoing shift from purely algorithmic black-box deep learning models towards the more explicit data modeling of classical statistics. Rather than viewing the modeling cultures as a dichotomy [35], our approach thus represents a step towards uniting the two worlds and combining their respective advantages [36].

Overall, our proposed strategy offers an integrated one-step modeling approach to extract activity signals from two-photon imaging data, operating directly on the video frames. Further, the approach exemplifies how tailoring a model to the specific properties of the data can provide an interpretable representation of the central patterns in the data, accounting for the distinct foreground and background activity as well as temporal correlations. It thus suggests that combining generative deep learning with structural information can also more generally be a promising approach for uncovering activity patterns in cell imaging data.

## 4 Materials and Methods

### 4.1 Typical steps during conventional bioimage analysis

The ultimate goal of bioimage analysis is to gain knowledge of biological processes by extracting relevant information from microscopy images, including time-lapse sequences that record the dynamics of the processes. In non-machine learning-based (‘conventional’) bioimage analysis, a sequence of operations is employed to extract the information [22, 23]. Depending on the specific application, this analysis workflow typically involves initial image processing (e.g., enhancement, noise reduction, filtering), followed by a sequence of operations that may include image segmentation and object detection (e.g., cell outline, nuclei), tracking of object movements, and intensity-based quantification and classification of objects (e.g., brightness, distance, size, co-localization). Finally, downstream data visualization and analytics as well as mathematical or statistical modeling are applied to allow a meaningful interpretation of the biological results, especially by comparing different experimental conditions. Depending on the complexity and size of the raw data set, it is necessary to design automated custom workflows that subsequently apply these operations to the data.

In conventional computational bioimage analysis, the main advantage of non-deep-learning approaches is that the underlying algorithms are transparent with observable input-output data, so that the results are readily interpretable. However, due to recent technical advances in microscopy methods and increased experimental complexities (e.g., in-vvol time-lapse imaging of the brain activity of behaving animals), the produced bioimage data has enormously increased in size and complexity. Conventional analysis tools are limited in processing these highly complex and large data sets, because meaningful results can only be obtained after tedious optimization of the algorithms. In addition, subtle differences in biological structures and dynamics are easily overseen or masked by suboptimal parameter settings that require individual manual adjustment. For example, existing toolbox solutions [14] account for the high complexity of the data with a large set of predefined parameters that require a deep knowledge of the underlying software by the user.

Driven by these challenges, a major paradigm shift has occurred with the massive adoption of deep learning technologies that are rapidly replacing conventional bioimage analysis approaches. Especially the use of artificial neural networks, and more recently of VAEs, in bioimage analysis offers considerable advantages for a flexible, yet efficient analysis of large, complex data sets.

### 4.2 Generative deep learning

As our proposed analysis strategy is based on a VAE, a generative deep learning approach first presented in [2], we first give a brief introduction to neural networks and generative deep learning in general, before presenting the VAE in more detail.

An *artificial neural network* (ANN) is a function composition 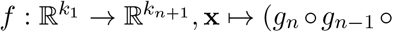 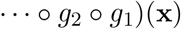 for distinct continuous functions 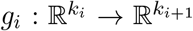 called the *layers* of the network. Each layer is of the form *g*_*i*_(**x**) = *h*_*i*_(**W**_*i*_**x**+**b**_*i*_), where *h*_*i*_ : ℝ → ℝ is a continuous non-linear function called the *activation function* and is applied element-wise, and 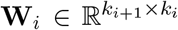 and 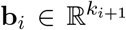 are *weights* and *biases*, also called *parameters*, of the network. Intuitively, by combining many of these layers of linear combinations followed by a non-linear activation into a *deep* network, ANNs can be used to model potentially very complex, non-linear structures in the data.

The process of finding a parameter set, i.e., determining weights and biases such that an ANN approximates a specific input-output mapping is called *learning* or *training* of the ANN. Thus, the term *deep learning* refers to the process of approximating potentially complex mappings with deep ANNs. More precisely, training is performed by defining a loss function as the training objective and repeatedly applying the chain rule to obtain partial derivatives of the loss with respect to the network parameters in order to minimize the loss function by stochastic gradient descent [24]. This training strategy results in the propagation of gradients ‘backwards’ through the network and is hence termed *backpropagation* [25]. ANNs can be trained to approximate various types of mappings from input data to outputs, e.g., mapping a dataset of images to binary classifications or segmentation masks, and have enjoyed great successes in many of these supervised tasks. They can also be employed as *generative* models that learn a joint distribution over all input variables in an unsupervised fashion to approximate the true underlying data distribution. Such a generative deep learning model then allows to draw synthetic samples from the learned distribution. Various approaches for such deep generative models have been proposed, including deep Boltzmann machines (DBMs) [26], generative adversarial networks (GANs) [27], and VAEs [2], which differ predominantly in the way the underlying data distribution is represented by the model.

### 4.3 Variational autoencoder

VAEs learn explicit parametrizations of the underlying probability distributions by employing two distinctly parametrized, but jointly optimized neural networks that are responsible for encoding and decoding of the data into and from a latent space that is governed by a probability distribution: The encoder maps the input data **x** to a lower-dimensional latent representation given by a random variable **z** by approximating the conditional distribution of **z** given **x**, while the decoder performs the reverse transformation from the latent space back to data space, parametrizing the conditional distribution of **x** given **z**.

The VAE training objective is to recover the central factors of variation underlying the data in the lower-dimensional latent space, thus obtaining a compressed representation based on which the input data distribution can be approximated. Since an ANN is used to encode the data into the latent space, the true posterior distribution *p*(**x|z**) becomes intractable. Hence, a variational approximation *q*(**z|x**) is employed – typically a Gaussian distribution with diagonal covariance matrix. A loss function for the model can then be derived based on variational inference [28]:

Minimizing the *Kullback-Leibler divergence* 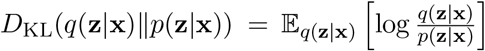 to the exact posterior is equivalent to maximizing a lower bound on the true data likelihood, the *evidence lower bound* (ELBO) given by 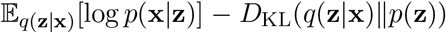. Denoting the parameters of the encoder and decoder neural networks with ***ϕ*** and ***θ***, respectively, we can define the VAE training objective as the negative ELBO:

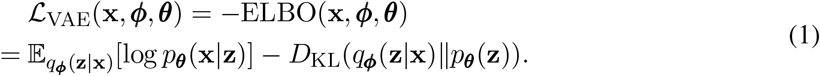

Intuitively, the first term in (1) can be thought of as a reconstruction error that encourages densities placing mass on configurations of latent variables that explain the observed data, while the second term has a regularizing effect by enforcing consistency between the prior and posterior of **z**. By maximizing the ELBO with respect to both ***θ*** and ***ϕ***, we can derive both approximate maximum likelihood estimates for ***θ*** and an optimal variational density *q*_*ϕ*_ [28].

In practice, with a Gaussian prior and posterior of **z**, the Kullback-Leibler divergence in (1) can be calculated analytically, while the expectation with respect to the variational posterior *q*_***ϕ***_(**z**|**x**) has to be approximated by Monte Carlo sampling. Using *S* samples for this approximation and a *K*-dimensional latent space the ELBO of a single data point **x**^(*i*)^ ∈ {**x**^(1)^, … , **x**^(*N*)^} can be calculated as

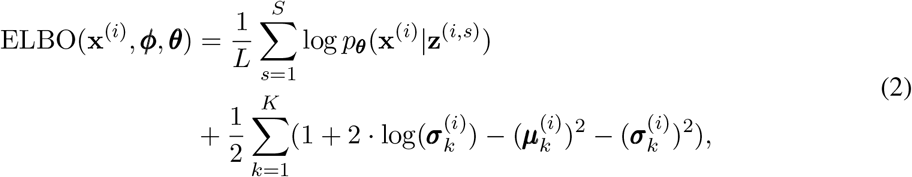

where ***μ*** ∈ ℝ^*K*^ and ***σ*** ∈ ℝ^*K*^ denote the parameters of the variational posterior (hence depending on its parameterization by ***ϕ***) and ***μ***_*k*_ denotes the *k*-th component of the vector. The loss over all data points **x**^(1)^, … , **x**^(*N*)^ is then given by

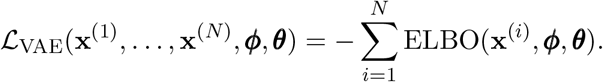

In our implementation, we use *S* = 1 throughout and follow the common practice of adding an 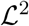 regularization term 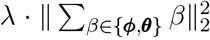 to prevent exploding model parameters. While obtaining an estimate of the gradient with respect to ***θ*** is straightforward, we have to employ a change of variables called the *reparameterization trick* [2] to estimate the gradient with respect to the variational parameters ***ϕ***. Intuitively, instead of sampling **z** ~ *q*_***ϕ***_(**z**|**x**), we sample some ***ϵ*** from a random variable independent of **x** and express **z** as a deterministic transformation of ***ϕ*** and ***ϵ***. Thus, we obtain unbiased estimates of the ELBO with respect to both ***θ*** and ***ϕ*** that are optimized with stochastic gradient descent [24].

### 4.4 Convolutional neural networks

For data in the form of two-dimensional arrays such as images, typically *convolutional neural networks* (CNN) are employed, a type of ANN specifically adapted to the spatial structure of image data [29]. CNNs comprise mainly two types of layers, namely *convolutional* layers and *pooling* layers.

A convolutional layer is defined by a number of filters, also called receptive fields, where each filter is represented by a weight matrix. The filters are applied locally, i.e., the weight matrix is multiplied element-wise with local patches of the input image and all values are summed up (mathematically, this corresponds to a discrete convolution, hence the name). Finally, a non-linear activation function is applied to the resulting weighted sum. We can thus think of a filter as a window sliding over the image, convolving the filter weight matrix with different patches of the image.

Each filter is defined by the size of the filter matrix, the stride, i.e., the step size with which it slides across the image, and its corresponding activation function. The locally applied convolution operation accommodates central characteristics of array data such as images, namely the fact that they often exhibit distinctive local motifs, representing highly correlated local groups of values, and that such local motifs can appear in any part of the image, i.e., are invariant to their overall position [29]. This is reflected in the CNN architecture by the idea of using the same weights at different locations, realized by the filter matrix applied across the image.

The convolutional layer outputs a feature map, which is subsequently aggregated in a pooling layer to form higher-level features and detect motifs by building more coarse-grained representations [29]. Technically, this is achieved by aggregating a local patch of a feature map to a single value, e.g., by computing the maximum or average of all values in the patch. Thus, a pooling layer is defined by the corresponding pooling function and the size and stride of the pooling filter. The parameters of the CNN are given by the set of all layer filter matrices and can be estimated with backpropagation as in ANNs.

### 4.5 Separating foreground and background: the dual-target VAE

Formally, we can describe the dataset **x** = {**x**^(1)^, … , **x**^(*N*)^} of images **x**^(*i*)^ ∈ ℝ^*L*×*M*^ for some *L, M* ∈ ℕ as a combination of foreground signal **x**_1_ and background signal **x**_2_, such that **x** = *f*(**x**_1_, **x**_2_) for some unspecified function *f* that describes the merging of foreground and background to the overall image.

Here, we consider an approximation of the background **x**_2_ by scaling down the data in order to minimize the bias introduced by the foreground signal **x**_1_, that has a much higher resolution. We choose *g* : ℝ^*L*×*M*^ → ℝ^*l*×*m*^ with *L* = *ls*, *M* = *ms* for some *l, m* ∈ ℕ to be a pooling function with stride *s*, that samples patches of size *s* down to a single pixel, thus creating a smaller image that averages out the fine-grained foreground signal, retaining mostly large-scale background activity.

As the foreground signal **x**_1_ cannot be approximated in a similarly simple fashion, we employ a pragmatic solution by introducing a cut-off and only retain pixel values larger than this threshold, corresponding to pixels with high activity. Thus, we approximate **x**_1_ by 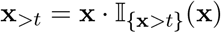.

To model foreground and background separately, we introduce separate VAE models with latent variables **z**_1_ and **z**_2_, that form compressed representations of **x**_1_ and **x**_2_ respectively. Accordingly, we consider likelihoods 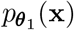 for the foreground and 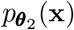 for the background and define corresponding variational encoders, 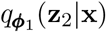 and 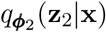 realized as CNNs.

To account for the different nature of sharp and local foreground signal and blurry, large-spanning background signal, we choose a small receptive field with a size similar to the regions of interest in the foreground for *q*_***ϕ***_(**z**_1_|**x**), while 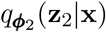 is parametrized by a CNN with larger receptive field. To obtain a joint model of the entire image including both foreground and background, we need to allow for interaction between the two models and thus add a fully connected ANN layer that integrates **z**_1_ and **z**_2_ into a joint latent space, before passing the output into the separate decoders. More specifically, the encoding and generative process of the resulting dual-target VAE model is given by

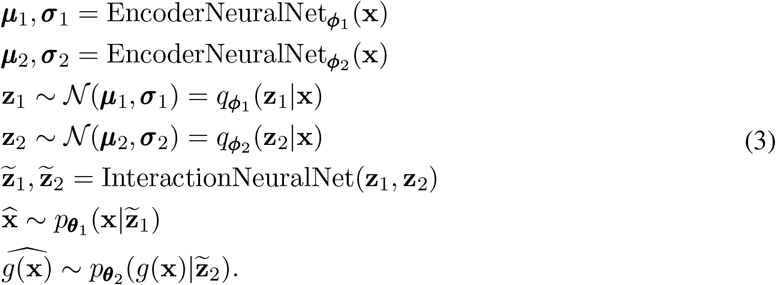

Consequently, we can derive an optimization criterion for the overall model by considering both ELBOs from the foreground and background part:

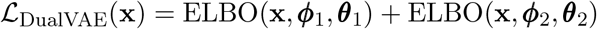

Estimators for the two ELBOs are calculated as in (2), such that the model can thus be trained by stochastic gradient descent analogously to the vanilla VAE (see Section 4.3). Note that the two coupled VAE models are not deterministic functions, as each includes drawing a sample from the latent variable **z** via a reparameterization (see Section 4.3). Also, because the latent spaces interact, each function includes the computation of both encoders.

The implementation of the approach and the training procedure of the dual-target VAE loss are illustrated and described in more detail in the accompanying Jupyter notebook.

### 4.6 Smoothing the latent representation with a kernel

To take into account the temporal correlation between subsequent frames, we additionally propose a smoothing step in the latent representation. Instead of training the model on randomly selected batches comprised of arbitrarily temporally distant frames, we consider neighborhoods of adjacent frames for training and thus make information about temporal proximity accessible to the model. For each neighborhood, we take a kernel-weighted average of the latent representations of all frames in the neighborhood, and use this smoothed latent representation as input for the decoder and loss function.

Formally, for some *m* ∈ ℕ, the neighborhood of size 2*m* + 1 of the *i*-th frame **x**^(*i*)^ is then given by *N*_*m*_(**x**^(*i*)^) ≔ (**x**^(*i*−*m*)^ **x**^(*i*−*m*+1)^ … **x**^(*i*)^ … **x**^(*i*+*m*)^). Denoting with *L* the sum of dimensions of the latent variables *z*_1,2_, a kernel function *K* : ℝ^*L*×(2*m*+1)^ → ℝ^*L*^ is applied to the latent representation mean *μ* = (*μ*_1_, *μ*_2_) for the respective posterior distributions of the two VAE models. As a kernel, we use the tricube function defined by *K* (*x*) = (1 − (|*x*|)^3^)^3^ and use the distance in frames to compute the weights with the kernel.

Instead of batch learning, we can now employ *neighborhood learning*: Instead of randomly partitioning the data set in disjoint batches ***B***_1_, … , ***B***_*J*_, and calculating the loss of the entire batch as 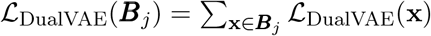 before applying one gradient update of the model parameters, we form a neighborhood ***N*** _*m*_(**x**) of size 2*m* + 1 around each **x** as defined above. With this, we can derive one loss value for the neighborhood by using the latent representation obtained as a weighted average of all frames in the batch with weights given by the kernel, i.e.,

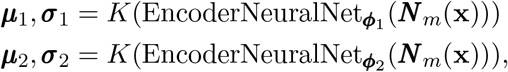

and subsequently use ***μ***_1_, ***σ***_1_, ***μ***_2_, ***σ***_2_ as in (3) to obtain samples **z**_1_, **z**_2_ and reconstructions, 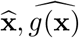. Thus, from each frame **x** and its respective neighborhood ***N*** _*m*_(**x**), we calculate a single value of 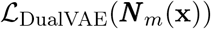 before applying one gradient update. Note how in this case the complexity for computing the gradients of each frame increases, as it has to flow through 2*n* additional computations of the EncoderNeuralNet.

To further explore our approach to obtain a smoother latent representation, we compare kernel smoothing in the latent space described above to smoothing performed at the level of the loss values of all frames in a neighborhood. This means that we calculate the loss 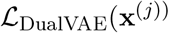 for a frame ***N*** _*m*_(**x**), for all **x**^(*j*)^ ∈ ***N*** _*m*_(**x**), and obtain a common loss as weighted average

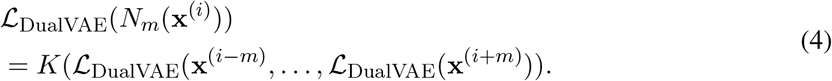

### 4.7 Data and preprocessing

We use data obtained from *in-vivo* two-photon calcium imaging of hippocampal CA1 neurons in mice. Mice were intra-hippocampally injected with adeno-associated viral constructs (AAV1.Syn.GCaMP6f. WPRE.SV4, University of Pennsylvania Vector Core) that established a panneuronal expression of the calcium-indicator GcaMP6f in CA1 neurons. Subsequently, a 3mm wide cranial window was implanted above the hippocampal formation to provide optical access to the structure. For details of the procedure, please see [11]. During the imaging mice were head fixed and placed on an air-floating styrofoam ball that allowed them to navigated through a virtual reality which was projected to four screens placed around them. Mice were trained to run along 4m long linear tracks to obtain goal-oriented rewards. Imaging was performed using a resonant/galvo high-speed laser scanning two-photon microscope (Neurolabware) with a frame rate of 30Hz for bidirectional scanning. The microscope was equipped with an electrically tunable, fast z-focusing lens (optotune, Edmund optics) to switch between z-planes within less than a millisecond. GCaMP6f was excited at 930nm with a femtosecond-pulsed two-photon laser (Mai Tai DeepSee, Spectra-Physics). To maximize the number of recorded neurons we scanned three imaging planes (≈ 25*μ*m spacing between planes) in rapid alternation so that each plane was sampled at 10Hz.

As a preprocessing step, we performed motion correction with the NoRMCorre algorithm [30] and subsequently normalized the data.

### 4.8 Implementation

The experiments have been implemented in the Julia programming language (version 1.4.1) and were carried out on a Linux cluster utilizing 150GB of RAM, 16 CPU-cores, and one NVIDIA Tesla V100 GPU. Model training was realized using the Julia machine learning library Flux.jl (version 10.1.0) and CUDA.jl (version 0.1.0) for GPU support.

To ensure comparability, all considered models (VAE with a CNN architecture, dual-target VAE with and without kernel smoothing) use the same architecture for their encoders and decoders (e.g., the encoder of the foreground model in the dual-target VAE is the same as the encoder of the convolutional baseline). In the following, we briefly describe the implementation of the different models. Details on the model structure and parameters are given in the supplementary material.

For the simple CNN-based VAE, we use 3 convolutional layers and one pooling layer for the CNN encoder and 4 transposed convolutional layers for the decoder. The first layer of the encoder is configured with a small filter of size 25 by 25. To keep the computational cost of the fully-connected ANN approach comparable to the CNN scenario, both encoder and decoder were realized as one-layer ANNs. In the dual-target VAE, the foreground model has the same architecture as the CNN-based VAE described above. For the background model, we employ of a two-layer convolutional encoder with a receptive field of size 100 by 100 and a fully-connected ANN decoder. The training target of the background VAE is the scaled version of the original data as described above in Section 4.5.

The training data comprises 10000 contiguous, normalized frames of size 796 by 512 pixels. The smaller target of size 8 by 5 for the background models is obtained by padding with zeroes and pooling 100 × 100 patches into their sums of pixels and normalizing subsequently. For pretraining, we choose *t* = 0.3 as threshold of the foreground approximation. We use the ADAM optimizer [31] for stochastic gradient descent and train the models for 20 epochs on the cut-off data and another 20 epochs on the original data with a batch size of 5. For the approaches with kernel smoothing, we use the parameters of the dual-target VAE as initialization and train for another 20 epochs and a neighbourhood of size 5.

The source code is available at https://gitlab.imbi.uni-freiburg.de/maren/incorporatingstructural-knowledge-into-unsupervised-deep-learning-for-two-photon-imaging-data

## Acknowledgements

The authors would like to thank Thomas Hainmueller, who generated the two-photon imaging data. Further, the authors acknowledge support by the state of Baden-Württemberg through bwHPC.

## Author contributions

Conceptualization: HB

Data curation: AK, MB

Methodology: HB, FE, MH, CB

Investigation: FE, MH, CB

Visualization: FE, MH

Writing – original draft: MH, FE, JS, AK, MB

Writing – review and editing: all authors

## Data and materials availability

Code to reproduce the figures in the manuscript and a Jupyter notebook tutorial illustrating the main functionalities of our method are available in a public Gitlab repository at https://gitlab.imbi.uni-freiburg.de/maren/incorporating-structural-knowledge-into-unsupervisedeep-learning-for-two-photon-imaging-data. In the repository, we also provide a small exemplary dataset to recapitulate our analyses.

