## Supplemental File 1 for "Incorporating structural knowledge into unsupervised deep learning for two-photon imaging data" (from another version of this preprint: version 1, 2021-05-18)

May 18, 2021

This document provides the supplementary tables with details about the model architectures mentioned in the manuscript.

### Model Architectures

#### Fully-connected Variational Autoencoder

| Encoder | | |
|---|---|---|
| Layer | Size | Activation |
| Input | $407552 = 796 \times 512 \times 1$ | |
| Dense | $240 = 10 \times 6 \times 4$ | leakyrelu |

| z-Layer and Interaction | | |
|---|---|---|
| Layer | Size | Activation |
| Dense ($\mu$) | 8 | identity |
| Dense ($\log \sigma^2$) | 8 | identity |
| Dense | $240 = 10 \times 6 \times 4$ | relu |

| Decoder | | |
|---|---|---|
| Layer | Size | Activation |
| Dense | $407552 = 796 \times 512 \times 1$ | leakyrelu |

#### Convolutional Variational Autoencoder

| Encoder | | |
|---|---|---|
| Layer | Size | Activation |
| Input | $796 \times 512 \times 1$ | |
| Conv | $130 \times 82 \times 16$ | leakyrelu |
| MaxPool | $43 \times 27 \times 16$ | |
| Conv | $20 \times 12 \times 16$ | leakyrelu |
| Conv | $10 \times 6 \times 4$ | leakyrelu |

| z-Layer and Interaction | | |
|---|---|---|
| Layer | Size | Activation |
| Dense ($\mu$) | 8 | identity |
| Dense ($\log \sigma^2$) | 8 | identity |
| Dense | $240 = 10 \times 6 \times 4$ | relu |

| Decoder | | |
|---|---|---|
| Layer | Size | Activation |
| ConvTranspose | $21 \times 13 \times 16$ | leakyrelu |
| ConvTranspose | $45 \times 29 \times 16$ | leakyrelu |
| ConvTranspose | $131 \times 83 \times 16$ | leakyrelu |
| ConvTranspose | $796 \times 512 \times 1$ | leakyrelu |

##### Dual-target Variational Autoencoder

| Encoder1 | | |
|---|---|---|
| Layer | Size | Activation |
| Input | $796 \times 512 \times 1$ | |
| Conv | $130 \times 82 \times 16$ | leakyrelu |
| MaxPool | $43 \times 27 \times 16$ | |
| Conv | $20 \times 12 \times 16$ | leakyrelu |
| Conv | $10 \times 6 \times 4$ | leakyrelu |

| Encoder2 | | |
|---|---|---|
| Layer | Size | Activation |
| Input | $796 \times 512 \times 1$ | |
| Conv | $30 \times 19 \times 4$ | leakyrelu |
| MaxPool | $8 \times 5 \times 4$ | |

| z1-Layer | | |
|---|---|---|
| Layer | Size | Activation |
| Dense ($\mu$) | 8 | identity |
| Dense ($\log \sigma^2$) | 8 | identity |

| z2-Layer | | |
|---|---|---|
| Layer | Size | Activation |
| Dense ($\mu$) | 2 | identity |
| Dense ($\log \sigma^2$) | 2 | identity |

| Interaction | | |
|---|---|---|
| Layer | Size | Activation |
| Dense | $240 + 160 = 10 \times 6 \times 4 + 8 \times 5 \times 4$ | relu |

| Decoder1 | | |
|---|---|---|
| Layer | Size | Activation |
| ConvTranspose | $21 \times 13 \times 16$ | leakyrelu |
| ConvTranspose | $45 \times 29 \times 16$ | leakyrelu |
| ConvTranspose | $131 \times 83 \times 16$ | leakyrelu |
| ConvTranspose | $796 \times 512 \times 1$ | leakyrelu |

| Decoder2 | | |
|---|---|---|
| Layer | Size | Activation |
| Dense | $40 = 8 \times 5 \times 1$ | sigmoid |
